# Assessment of a Passive Sampler for Aquifer Microbial Community Profiling and Comparison of Porous Media Applied in a Managed Recharge Scenario

**DOI:** 10.64898/2025.12.19.695546

**Authors:** Mia Riddley, Haeil Byeon, Keegan Trubenbach, Madeline Schreiber, Amy Pruden, Jingqiu Liao

## Abstract

Microbial communities play an important role in aquifer systems, as both hazards (e.g., pathogens, carriers of antibiotic resistance genes) and contributors to contaminant remediation. However, monitoring native microbial communities *in situ* remains challenging. To address this challenge, we developed a passive sampler with that employs removable cartridges containing solid media designed to re-create the aquifer environment in a controlled fashion and support consistent, repeatable, time-series sampling. With a bench-scale, continuous-loop setup circulating advanced-treated wastewater, we compared microbial community dynamics across three candidate porous media: native aquifer sediment, zirconia beads, and laboratory-grade silica sand. 16S rRNA gene amplicon sequencing revealed that native aquifer sediment best reflected influent microbial composition and temporal shifts. Native aquifer sediment also exhibited more spatial consistency in microbial diversity along the sampler. These findings suggest that native aquifer sediment is an optimal porous medium for long-term, *in situ* monitoring of microbial communities for managed aquifer recharge and other applications in groundwater systems.

## 1. Introduction

Groundwater is an essential resource for drinking water, agriculture, and industry, especially in times of drought or surface water scarcity. Globally, approximately 69% of groundwater abstractions are used for agriculture, 22% for domestic purposes, and 9% for industry, underscoring groundwater’s critical role in supporting food production, water supply, and economic development (Pointet, 2022). However, groundwater systems (aquifers) are increasingly stressed by environmental pressures, particularly in coastal and urban regions (Mieno et al., 2024). One of the most pressing issues is seawater intrusion, driven by over usage of groundwater and sea level rise (Barlow & Reichard, 2010; Werner et al., 2013). Seawater intrusion reduces the availability of potable water and alters the geochemical conditions within aquifers. Additionally, land subsidence due to groundwater depletion affects infrastructure and accelerates saltwater encroachment (Davydzenka et al., 2024). Other environmental stressors include variability in precipitation, drought, mismanagement, climate change, and increasing demand from agriculture and urbanization, leading to long-term imbalances between recharge and withdrawal (Taylor et al., 2013). These complex stressors necessitate adaptive recharge strategies to ensure sustainable groundwater management.

Managed aquifer recharge (MAR) technologies offer a promising solution to mitigate the ongoing depletion of groundwater resources by injecting treated wastewater into an aquifer (Drewes, 2009). As of 2024, there are over 1200 MAR sites worldwide, with 31.7% utilizing well injection methods (IGRAC, 2025). While MAR plays a critical role in addressing future concerns regarding water storage and water quality, the use of reclaimed water in MAR introduces additional risk considerations, as it may contain residual contaminants, such as microbial pathogens, antibiotic resistance genes (ARGs), and persistent organic compounds (e.g., PFAS, pharmaceuticals). These constituents may persist in the effluent of treated wastewater, particularly if they are recalcitrant or difficult to biodegrade (Hong et al., 2013; Khan et al., 2022; Pruden, 2014). Such concerns underscore the potential value of monitoring aquifer microbial communities not only as an indicator of water quality, but also for understanding potential impacts of MAR on native microbial ecosystems and biogeochemical function (Blair, Garner, et al., 2024; Blair, Vaidya, et al., 2024). However, traditional monitoring of MAR systems primarily emphasizes chemical and hydraulic parameters (H. Zhang et al., 2020). Also, among 138 MAR-related risk assessment studies conducted globally, only 5% were carried out in the United States (Imig et al., 2022), highlighting a significant knowledge gap regarding microbial risks of MAR systems across a range of geographies.

Understanding microbial risks in MAR remains challenging because groundwater sampling typically requires drilling and constructing monitoring wells. These activities can perturb aquifer conditions and introduce artifacts or contaminants, thereby biasing the samples collected. One alternative is to construct laboratory-scale columns to simulate the aquifer system in the laboratory. Specifically, columns can be constructed at laboratory-scale to simulate the filtration and biogeochemical processes that occur as water infiltrates through soil into an aquifer (Sharma & Kennedy, 2017). Ideally, corresponding microbial ecological studies should capture all interfaces of the sampling environment, which in the case of laboratory columns would be the bulk effluent water and the column media over time (Drennan et al., 2017). However, long-term column studies restrict the removal of sediment with time because it entails opening the column and disrupting the flow conditions. Equipping field and lab studies with passive samplers provides an alternative means to collect microbial samples associated with solid media with time, without disturbing the sampling environment. This can enable time-series sampling needed to investigate dynamics with time and can be compared alongside sediment samples to obtain a fuller picture of the microbial ecology of the system. Commercially-available passive microbial samplers like the BIO-TRAP® have been used in groundwater remediation studies (Busch-Harris et al., 2008). These samplers use high surface area substrates (e.g., polymer or activated carbon matrices) to facilitate microbial capture and colonization. While effective for targeted bioremediation assessment, commercial samplers are limited in flow-dependency and not suited for broader community profiling, especially in MAR systems that may require distinct flow-specific configurations during their recharge processes. Alternatively, studies in aquatic environments have been carried out using in-house apparatuses, employing materials such as glass beads, cellulose, and artificial substrates to sample microbial biofilms or collect ambient microbial cells (Beak et al., 1973; Rickwood & Carr, 2009). While such approaches provide promising alternatives to commercial samplers, there is a lack of consensus and comparative studies on the optimal design for passive microbial samplers or the porous material used, especially when the goal is to capture microbial community composition that is representative of the aquifer and not altered by the sampler itself.

In the present study, we evaluated three candidate porous media, native aquifer sediment, zirconia beads, and laboratory-grade silica sand, used in passive microbial samplers within a controlled, bench-scale testing environment. We compared the effectiveness of the three media types in capturing microbial communities over time while being exposed to advanced treated wastewater used as recharge for an MAR project in Virginia. Aquifer sediment, being the host material for an MAR project, the Sustainable Water Initiative for Tomorrow (SWIFT) Center, located in Suffolk, Virginia, was hypothesized to support microbial colonization patterns that best reflect the *in-situ* community. Zirconia beads, widely used in DNA extraction protocols for cell disruption and DNA binding, were selected as a potentially effective high-capture medium that maximizes DNA recovery (Epperson & Strong, 2020). Laboratory-grade silica sand is widely used in drinking water treatment plants for its effectiveness in removing turbidity and microbes and was selected as a relatively low biomass and permeable substrate (Maiyo et al., 2023). We evaluated the media using 16S rRNA gene amplicon sequencing and a series of statistical tests. Overall, the present study highlights the practical importance of selecting an optimal porous medium for the further development of full-scale passive microbial samplers to monitor microbial processes in MAR systems.

## 2. Methodology

### 2.1 Experimental setup and porous media

A total of 20, 4”x1” hollow cylindrical columns were designed in SOLIDWORKS 2023-2024 and printed using Overture PLA 1.75 mm filament on a Intamsys Funmat HT printer. Columns were designed for a screw fitting with Bluefin 1/4" Hose Barb x 1/2" NPT Male Brass Pipe Adapter fittings. ¼” I.D. x ⅜” O.D. UDP clear vinyl tubing was connected to the hose barb and set into the Cole-Parmer Instrument Co. 7520-10 Masterflex L/S Variable Speed Drive peristaltic pumps system (**Figure S1**).

Three porous media were assessed in this study: native aquifer sediment, zirconia beads, and laboratory grade sand. Aquifer sediment consisted of well cuttings (sediment + drilling mud) from a depth interval of 657.5 - 806.5 ft below ground surface that were collected during drilling of the Potomac Aquifer System for installation of MAR recharge wells. Prior to use in the experiments, the sediment was washed onsite using the following methods. First, the sediment-mud mix was washed 3 gallons at a time within a rotating mixer drum. The wet media was then sieved through a #270 screen, and the process was repeated three additional times. Next, sediment was sieved through a #4 screen and laid out flat to dry over tarp, outdoors, under ambient conditions. Drying sediment was sheltered under a tent to decrease the influence of weather conditions. After drying, the particle size was measured using sieve analysis and was characterized as sand. Upon arrival in the lab, the sediment was rinsed in a 70% ethanol bath and was air-dried for 24 hours. Following drying the sediment was added into a sterile 1L Nalgene bottle, capped, and autoclaved at a Gravity 1 setting. The sediment was then laid out to dry for 24 hours, and this process was repeated three times. White, fine, laboratory-grade sand (Carolina, Burlington, NC) and MSE PRO 0.1 mm Zirconia Beads for DNA, RNA and Protein Extraction and Isolation (MSE Supplies, Tucson, AZ) were cleaned in repeated manner.

A continuous loop system was used to test the capture behavior of all three porous media. Influent reservoirs for the continuous loop systems consisted of HDX 20-gallon totes (Home Depot, Atlanta, GA) containing 16 liters of advanced treated wastewater collected from the SWIFT facility. The water was cycled through the media columns (samplers) and the influent reservoirs at a rate of 10 mL/min over a 21-day period. To achieve experimental replication, three parallel columns were run for each media type. All porous media were autoclaved prior to loading into each column. Separately, mesh squares, fittings, tubes, and 3D printed columns were soaked in 70% ethanol for 24 hours prior to assembly. Upon setting up, 1L of 70% ethanol was cycled through the system and totes were disinfected using a 70% ethanol wash and followed with a bleach wash. All materials were allowed to dry prior to the loading of porous media.

### 2.2. Sampling of porous media and influent reservoirs

Samples were collected in triplicate every 7 days (Day 7, 14, and 21) from both the top 1/4^th^ of the sampler and the bottom 1/4^th^ of the sampler of each column containing media, as well as from their respective influent reservoirs. An initial sample was also taken from each media type and the influent water at the start of the experiment, yielding a total of 67 samples over the 21-day experiment period. Samples of approximately 1-g of media were collected from both the top and bottom of each column using a sterile spatula. Media samples were transferred to 15 mL microcentrifuge tubes and stored at –80 °C until DNA extraction. Influent reservoir samples were taken as 1-L water from the reservoir tote. Immediately after collection, the influent reservoir samples were filtered through a vacuum pump over a 0.45-uM filter (MilliporeSigma, Burlington, MA). Flow through was discarded and filters were stored in a 2-mL microcentrifuge tube at -80°C and filters were shredded using sterile tweezers for each influent reservoir sample prior to DNA extraction.

### 2.3. DNA extraction and 16S rRNA gene amplicon sequencing

DNA was extracted from 0.5-g of each media sample and shredded filter of each influent reservoir water sample using the FastDNA Spin Kit for Soil (MP Biomedicals, Solon, OH). Polymerase chain reaction (PCR) was performed to amplify the V4 region of the 16S rRNA gene using 515F and 806R primers (5’ - GTG CCA GCM GCC GCG GTA A - 3’ and 5’ - GGA CTA CHV GGG TWT CTA AT - 3, respectively). PCR amplicons were confirmed by gel electrophoresis and further assessed for quality (*A*260/280 ∼1.80 and *A*260/230 ∼2.0) using NanoDrop (Thermo Fisher Scientific, Asheville, NC) and for quantity using a Qubit dsDNA BR kit (Thermo Fisher Scientific, Asheville, NC). PCR was completed in duplicates and pooled for each sample. The DNA libraries were constructed using Nextera XT DNA Library Preparation Kit (Illumina, CA, USA). DNA concentration normalization, barcoding, and sequencing were subsequently executed by the Genomics Facility at Cornell University using the MiSeq Illumina platform (V2 500 bp; paired-end).

### 2.4. Microbial community characterization and phylogenetic tree construction

Raw sequencing reads were pre-processed following our previously used methods based on the QIIME2 workflow (Liao, 2021). In brief, raw sequencing reads were denoised using DADA2, yielding a range of reads from 117 to 17982 among samples. Sequences with a similarity > 0.97 were clustered *de novo* into operational taxonomic units (OTUs) using q2-vsearch. OTUs were taxonomically classified by comparing sequences against the pre-trained classifiers Greengenes 13_8 using classify-sklearn. Bacterial OTUs with a frequency less than 2 were filtered out. After data preprocessing and filtering, a total of 1790 OTUs were generated. With these OTUs, four α-diversity metrics (richness, Shannon-Wiener diversity index, Simpson’s evenness, and Faith’s phylogenetic diversity [PD]), which refers to the diversity of organisms within a single sample, was calculated using QIIME2 q2-diversity plugin. Four β-diversity metrics (Bray-Curtis, Jaccard, unweighted UniFrac, and weighted UniFrac distances), which are used to compare the variability in community composition between samples, were also calculated using QIIME2 q2-diversity plugin. Additionally, OTUs were collapsed to each taxonomic rank (i.e., phylum, class, order, family, genus, and species) based on annotation.

Using the masked sequence alignment for representative sequences for OTUs, a phylogenetic tree was constructed using IQ-TREE with 1000 bootstraps (Minh et al., 2020). The best evolutionary model was determined based on the Bayesian information criterion (BIC) by the ModelFinder Plus implemented in IQ-TREE. The tree was rooted by mid-point and was visualized using iTOL (Letunic & Bork, 2024).

### 2.5. Statistical analysis

Pairwise correlations between α-diversity metrics and between β-diversity metrics were assessed using Spearman’s rank correlation tests and Mantel tests, respectively. α-diversity indicated by Shannon-Wiener diversity index as well as relative abundance of each taxon at each taxonomic rank was compared among three porous media i) overall, ii) at each sampling time point, iii) at each sampling location (top or bottom), and iv) among sampling time points and v) among sampling locations for each sampler medium using a Kruskal-Wallis test each followed by *post hoc* two-sided Mann-Whitney *U* tests. Principal Coordinate Analysis (PCoA) followed by a Permutational Multivariate Analysis of Variance (PERMANOVA) test was conducted to compare the dissimilarity in microbial community composition indicated by Bray-Curtis distance among the three porous media i) overall, ii) at each sampling time point, iii) at each sampling location (top or bottom of a sampler column), and iv) among sampling time points and v) among sampling locations for each sampler medium. False Discovery Rate (FDR) correction was applied to account for multiple testing. Statistical significance was defined as *P* < 0.05 and adjusted *P* < 0.1.

## 3. Results

### 3.1. Variation in microbial α-diversity across porous media was accentuated with time, with influent α-diversity best reflected by native aquifer sediment

Understanding microbial α-diversity is important for assessing ecosystem stability and water quality and helps identify shifts in microbial communities that may indicate environmental stress or changes. In this experiment, α-diversity also served to assess which porous media captured the most different kinds of microbes from a taxonomic standpoint. Microbial α-diversity was characterized using four indices, richness, Shannon-Wiener diversity index, Simpson’s evenness, and Faith’s PD. Richness was normally distributed, ranging from 7 to 191 with a mean of 88 (Figure S2a). Shannon-Wiener diversity index and Simpson’s evenness distributions were left skewed, ranging from 1.90 to 5.09 and 0.57 to 0.96, with a mean of 3.74 and 0.84, respectively (Figure S2a). Faith’s PD produced a slight right skew distribution, ranging from 1.47 to 5.74 with a mean of 3.42 (Figure S2a). Since Shannon-Wiener diversity index had the overall strongest correlations with other α-diversity indices (Spearman ρ > 0.6, *P* < 0.05 for all; Figure S2b), it was chosen as the representative index for subsequent microbial α-diversity comparisons across the experiment.

To understand the influence of porous media on microbial α-diversity, we assessed the differences in Shannon-Wiener diversity compared among sand, aquifer sediment, and zirconia beads. Results show that there was no significant difference compared among these porous media (Kruskal-Wallis *P* = 0.297, Figure S3). Based on the medians, aquifer sediment captured microbial α-diversity more similar to that of the influent at Day 0, which was characterized by a Shannon-Wiener diversity index of 3.96 (Figure S3). Stratifying by sampling day, significant differences in Shannon-Wiener diversity across porous media were observed on Day 21 (Kruskal-Wallis *P* = 0.042), with sand having a significantly lower microbial diversity compared to zirconia and aquifer sediment (adjusted Mann-Whitney *U P =* 0.042 and 0.038, respectively; **Figure 1a**). Stratifying by sampling location, no significant differences in Shannon-Wiener diversity was observed (Kruskal-Wallis *P* > 0.05 for all; **Figure 1b**). Across sampling days and locations, the α-diversity of the influent at Day 0 was generally more similar to that of the aquifer sediment. These results suggest that aquifer sediment most closely reflected microbial α-diversity of the corresponding aqueous influent. Also, microbial α-diversity diverged among porous media to varying degrees as time progressed.

**Figure 1.**
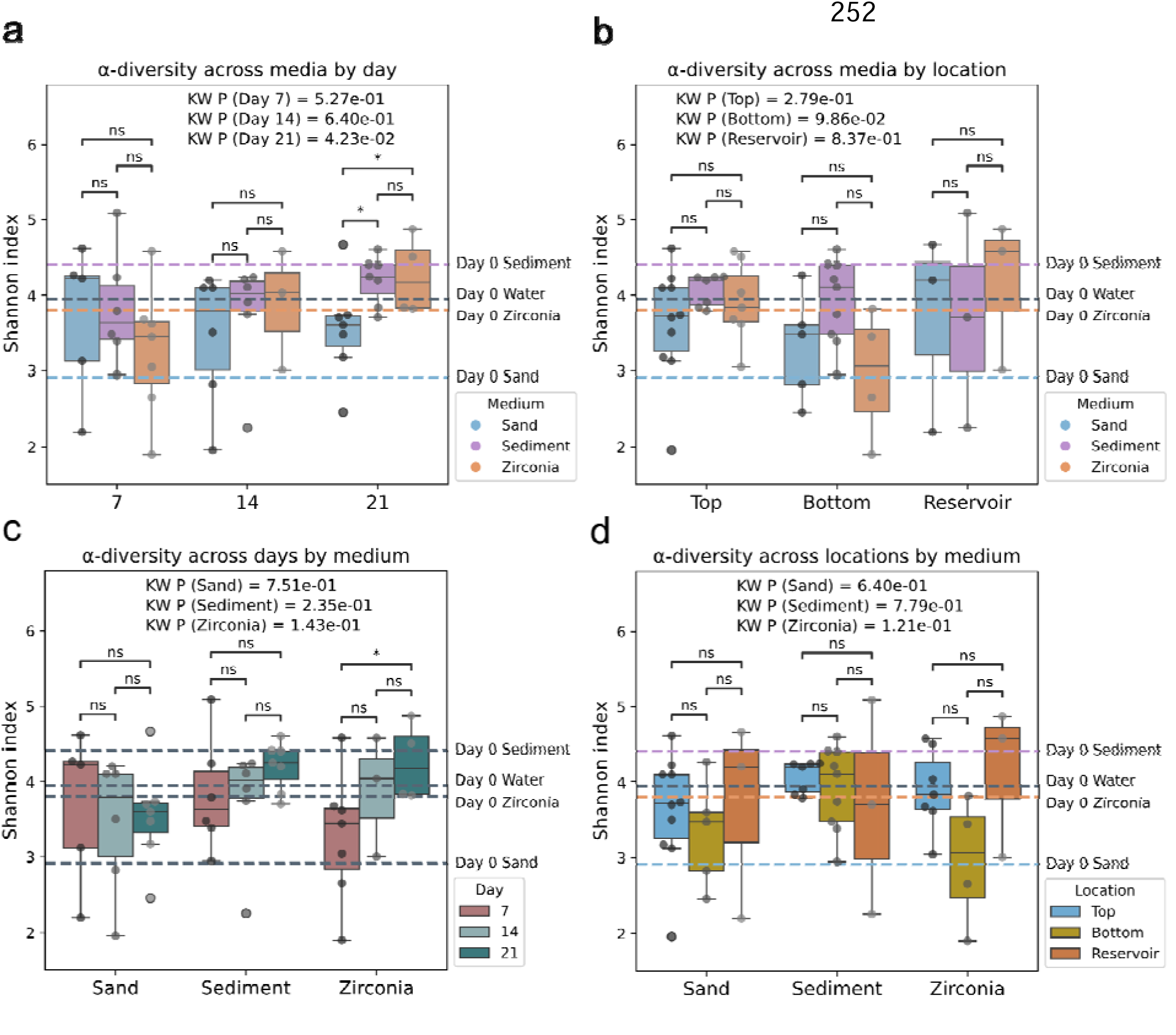
Microbial α-diversity captured on different porous media across sampling locations and days. Boxplots compare Shannon-Wiener diversity indices among (a) porous media (sand, aquifer sediment, and zirconia beads) at each sampling day (day 7, 14, or 21), (b) porous media at each sampling location (top, bottom of a sampler column, or influent reservoir), (c) sampling days for each medium, and (d) sampling locations for each medium. KW: Kruskal-Wallis. * And ns denotes two-sided Mann-Whitney U P < 0.05 and > 0.05 (not significant), respectively. Grouped box plots show the interquartile range (IQR), with the line representing the median and whiskers extending to 1.5 times the IQR.

To further understand the influence of sampling days and locations on microbial α-diversity within the sampler column, we assessed the differences in Shannon-Wiener diversity compared among Day 7, 14, and 21 as well as among the top and the bottom of the sampler column and influent reservoir for each medium. Results showed that Shannon-Wiener diversity of sand decreased over time, while that of aquifer sediment and zirconia beads increased (**Figure 1c**). The temporal changes in Shannon-Wiener diversity were not significant for sand and aquifer sediment, however (Kruskal-Wallis *P* = 0.751 and 0.235, respectively), while there was a significant difference in zirconia beads samples taken on Day 21 compared to Day 7 (Mann-Whitney *U P* = 0.042; **Figure 1c**). When comparing among sampling locations, while Shannon-Wiener diversity was not significant regardless of porous media (Kruskal-Wallis *P* = 0.640, 0.779, and 0.121 for sand, aquifer sediment, and zirconia beads, respectively), aquifer sediment exhibited considerably less variation among the top and bottom of the sampler column and the influent reservoir (**Figure 1d**). Overall, these results suggest that sampling time plays a critical role in monitoring microbial α-diversity with passive samplers, especially when using zirconia beads as the porous media, while the sampling location on the sampler has a limited impact on diversity, particularly when aquifer sediment was used.

### 3.2. Native aquifer sediment microbial communities best reflected temporal shifts in influent reservoir microbiota and were least affected by spatial variation in the sampler

In MAR systems, understanding microbial β-diversity - the variation in microbial community composition between samples - can provide insights into the acclimation of microbial communities to their sampling environment. In the case of this experiment, we aimed to observe the acclimation of the influent microbial community to each corresponding media with time. Microbial β-diversity was characterized using four indices, Bray-Curtis, Jaccard, unweighted UniFrac, and weighted UniFrac distances. Bray-Curtis, Jaccard, and unweighted UniFrac all produced a left skew distribution, ranging from 0.15 to 1, 0.44 to 1, 0.03 to 0.39 with a mean of 0.75, 0.85, and 0.23, respectively (**Figure S4a**). Weighted UniFrac produced a normal distribution ranging from 0.03 to 0.39, with an average of 0.23 (**Figure S4a**). Since Bray-Curtis distance had the overall strongest correlations with other β-diversity indices (Mentel ρ > 0.70, *P* < 0.05 for all; **Figure S4b**), it was chosen as the representative index of subsequent microbial β-diversity comparisons across the experiment.

To understand the influence of porous media on microbial β-diversity, we assessed the differences in Bray-Curtis distance compared among sand, aquifer sediment, and zirconia beads. PCoA showed that samples formed significant clustering by porous media (PERMANOVA *P =* 0.007, **Figure S5**), indicating that there were distinctions in the microbial community composition selected for and captured by different porous media. Stratifying by sampling day, microbial community composition did not differ significantly among porous media on Day 7 (PERMANOVA *P* = 0.064), but showed significant differences by Days 14 and 21 (*P* = 0.020 and 0.007, respectively; **Figure 2a**). In addition to sampling day, sampling location also made a difference in the dissimilarity of microbial community composition across porous media. As it pertains to the microbial communities identified at the same sampling location across the different media, specifically for the top and bottom samples. The effect was significant at the top and bottom of the sampler (*P* = 0.008 and 0.012, respectively), while no significant differences were observed across the three influent reservoirs of the media (*P* = 0.683; Figure 2b).

**Figure 2.**
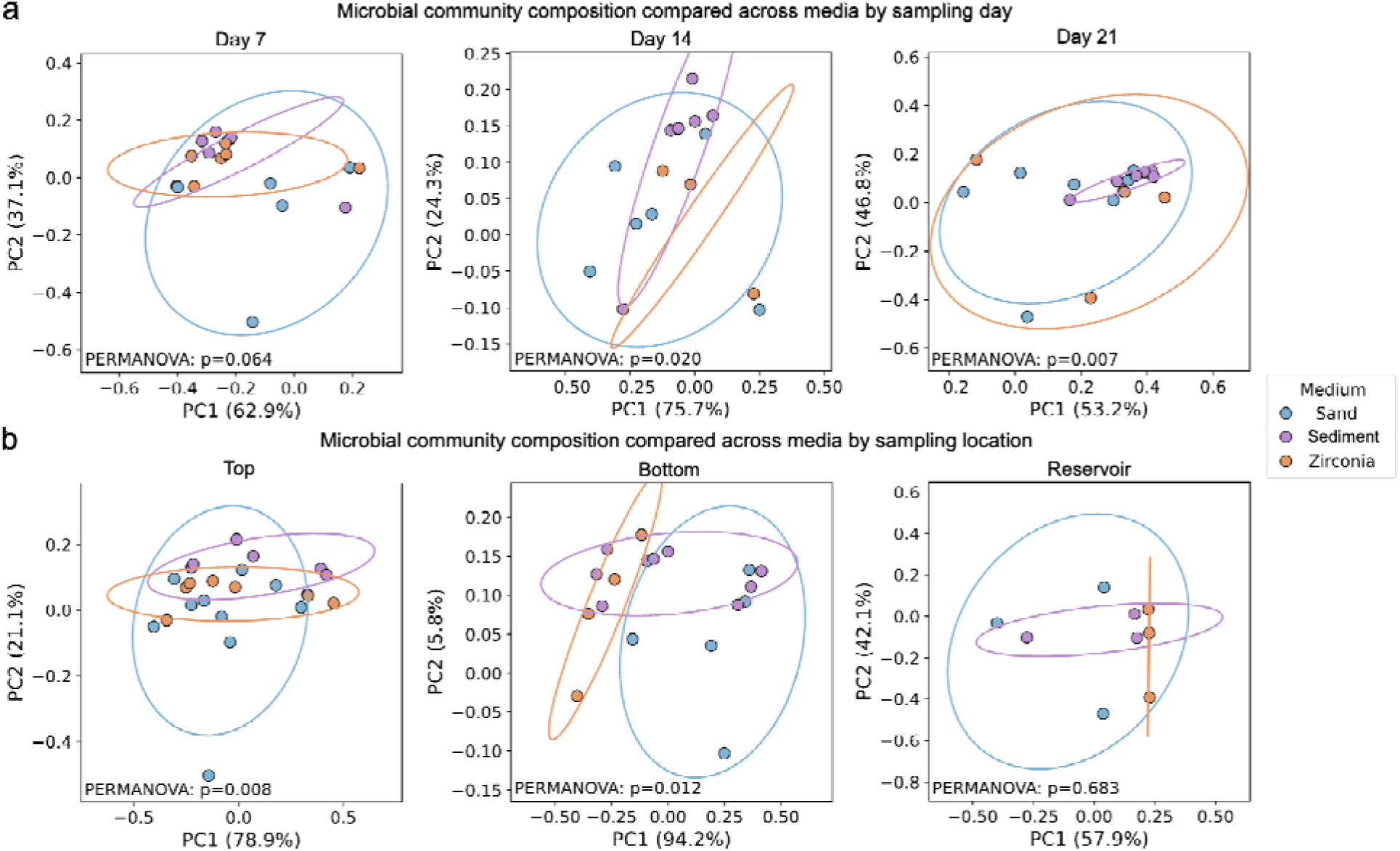
Microbial β-diversity captured on different porous media across sampling days and locations. PCoA plot showing clustering of samples based on Bray-Curti distance grouped by (a) porous media (sand, aquifer sediment, and zirconia beads) at each sampling day (day 7, 14, or 21), (b) porous media at each sampling location (top, bottom of a sampler column, or influent reservoir). PC1 (Principal Component 1) and PC2 (Principal Component 2) is the first and second principal coordinate, respectively. The percentage indicates the proportion of the total variance in the data that is captured by that specific axis. PERMANOVA: Permutational Multivariate Analysis of Variance.

To further evaluate the influence of sampling days and locations on microbial β-diversity for a given porous medium, we assessed the dissimilarities of community composition based on Bray-Curtis distances compared among Day 7, 14, and 21 as well as among the top and the bottom of the sampler column and influent reservoir for each medium. PCoA along with PERMANOVA showed that microbial community composition significantly shifted over time for all three porous media (PERMANOVA *P* < 0.05 for all; **Figure 3a**). Aquifer sediment exhibited the strongest microbial succession over time (*P* = 0.001), followed by zirconia (*P* = 0.005) and sand (*P* = 0.012). In addition, microbial community composition was associated with sampling locations for samplers containing sand or zirconia beads (*P* = 0.031 and 0.020, respectively), suggesting that there is a need to homogenize the samples taken from the top and bottom of the sampler when using these two porous media (**Figure 3b**). By contrast, microbial communities from aquifer sediment did not significantly differ in composition across the three sampling locations (top and bottom of a column, and an influent reservoir) (*P* = 0.259). These results suggest that among the tested porous media, aquifer sediment best reflected temporal changes in microbial β-diversity of the aqueous influent and was minimally affected by sampling location within the sampler, reducing potential need to homogenize samples coming from multiple depths.

**Figure 3.**
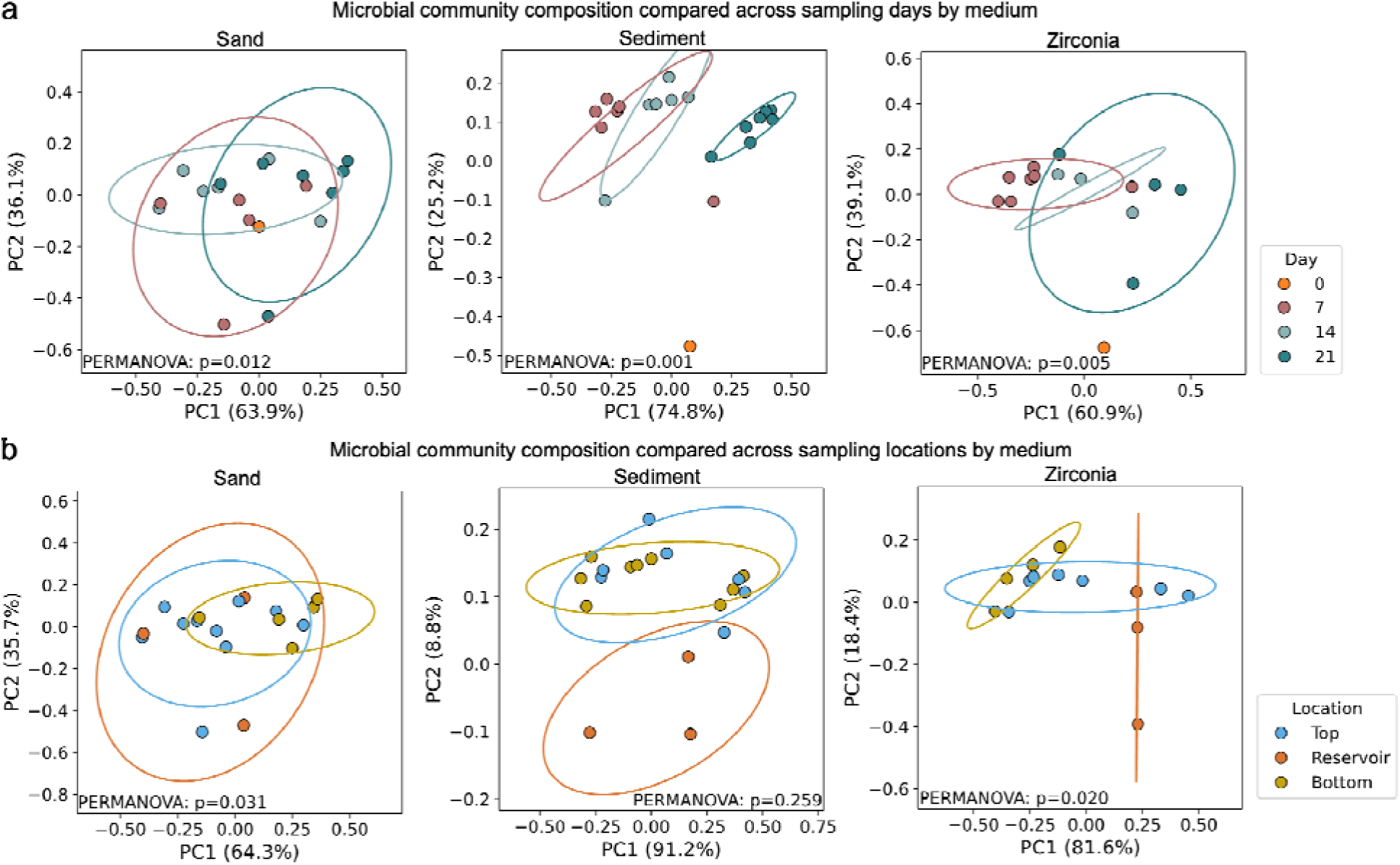
Microbial β-diversity captured at sampling days and locations across different porous media. PCoA plot showing clustering of samples based on Bray-Curtis distance grouped by (a) sampling days (day 0, 7, 14, or 21) and (b) sampling locations (top, bottom of a sampler column, or influent reservoir) for each medium (sand, aquifer sediment, and zirconia beads), respectively.

### 3.3. Proteobacteria, Bacteroidetes, Planctomycetes, and Verrucomicrobia in the aquifer sediment sampler exhibited sensitivity to temporal changes

As various microbial taxa drive critical processes, like biodegradation of trace organic carbon and nutrient cycling, investigating which microbial taxa persist and how they adapt to the sampler environment can support the design and management of engineered systems that promote beneficial microbial functions and long-term resilience. In this study, a total of 1,790 OTUs were identified, representing 20 microbial phyla (**Figure 4a**). Across all porous media, Proteobacteria and Bacteroidetes dominated at the phylum level (**Figure 4b**). Compared to media samples, corresponding influent reservoir samples for each porous media displayed a more diverse composition of prevalent phyla, which was most evident among influent reservoir samples for aquifer sediment at Day 7 and for sand and zirconia beads at Day 21 (**Figure 4b**). Betaproteobacteria and Gammaproteobacteria were the leading classes; Betaproteobacteria increased over time and were enriched in influent reservoirs, while Gammaproteobacteria were enriched at early timepoints and samples taken from the top of the sampler (**Figure S6**). Comamonadaceae and Rhodobacteraceae dominated at the family level. Comamonadaceae were consistently abundant at early timepoints and in upper sampler regions across all porous media types, while Rhodobacteraceae were enriched in influent reservoirs, especially at later timepoints (**Figure S7**). At the genus level, *Rhodoferax and Comamonas* dominated the sand and zirconia bead samples was prominent, particularly at early timepoints, whereas in aquifer sediment samples, *Rhodoferax and Paracoccus* were the most abundant, especially in transition zones within the sampler (**Figure S8**). These results highlight the key microbial taxa that passive samplers may capture from advanced treated wastewater used in MAR.

**Figure 4.**
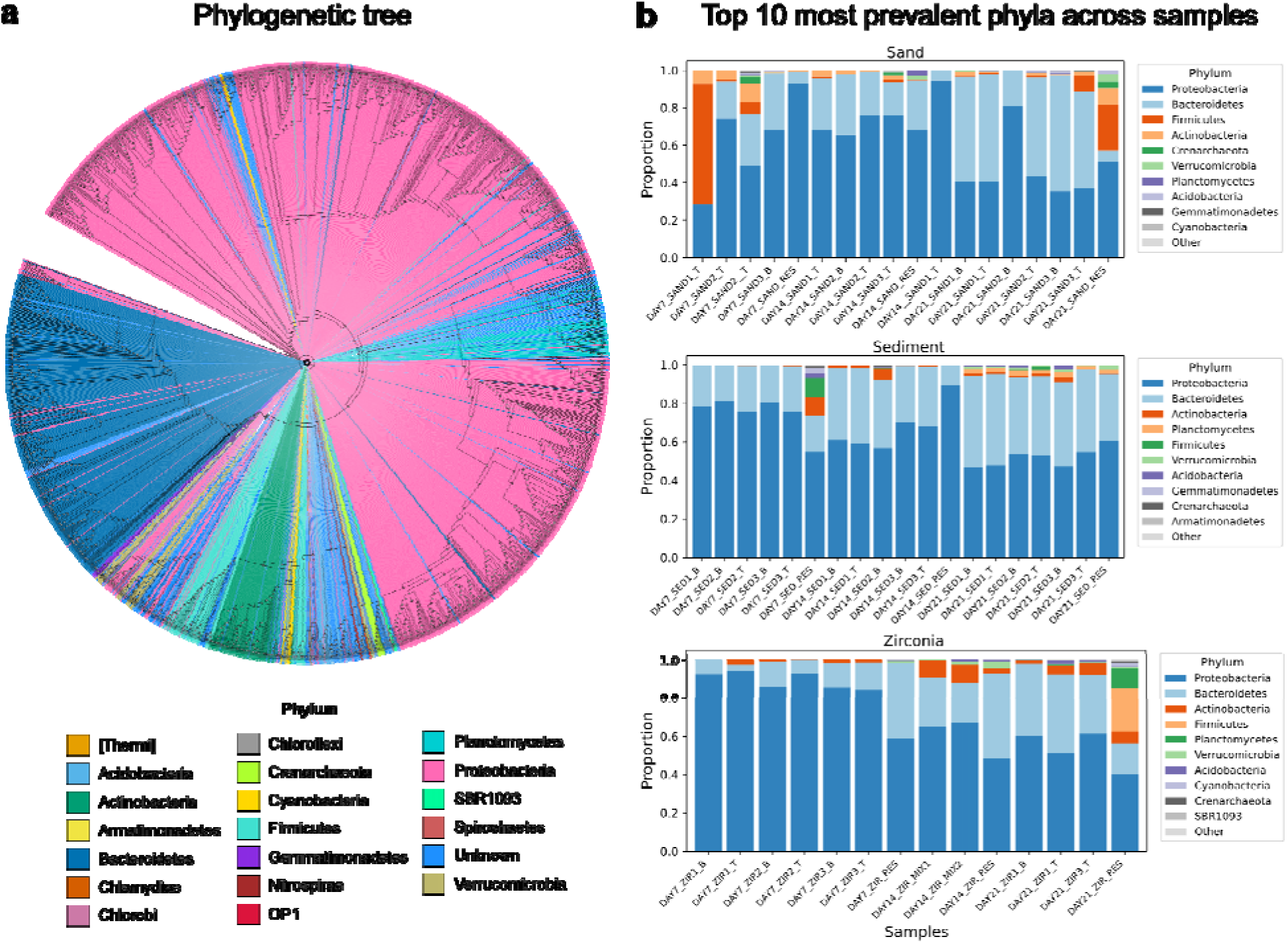
Phylogenetic relationships and taxonomic composition of microbial communities recovered by the porous media. (a) Phylogenetic tree of 1790 OTUs based on 16S rRNA gene sequences. The tree was constructed with 1000 bootstraps and rooted by midpoint. The branches are color-coded by the phylum that each OTU represents. (b) Stacked bar plots showing the relative abundance of the top 10 most dominant phyla across samples of each porous medium (sand [SAND], aquifer sediment [SED], and zirconia beads [ZIR]) from Day 7, 14, and 21. Abbreviations on the x-axis labels indicate bottom (B) and top (T) of a sampler column and influent reservoir (RES), respectively.

To further evaluate which microbial taxa were distinctly associated with the corresponding porous media, sampling day, and sampling location, relative abundances of each taxon were compared across Day 7, 14, and 21, and among the top and the bottom of each sampler column and across the three influent reservoirs for each porous media at each taxonomic rank. At the phylum level, Proteobacteria, Bacteroidetes, Plantectomycetes, and Verrucomicrobia had significantly different relative abundances among sampling days in aquifer sediment (Kruskal-Wallis *P <* 0.05 and adjusted *P* < 0.1 for all; **Figure 5**). Proteobacteria showed a significant decline in relative abundance over time, while Bacteroidetes, Planctomycetes, and Verrucomicrobia increased significantly by Day 21 relative to Day 7 (Mann-Whitney *U P* < 0.05 for all; **Figure 5**). In addition, five microbial classes (e.g., Gammaproteobacteria, Flavobacteria, and Planctomycetia), two orders (Ellin329 and Pirellulales), one family (Pirellulaceae), one genus (A17), and one Planctomycetes species exhibited significant differences in relative abundance in aquifer sediment across sampling days (Kruskal-Wallis *P <* 0.05 and adjusted *P* < 0.1 for all; **Table S1**). Also, one microbial class (i.e., Pedosphaerae) and one order (i.e., Opitutales) showed significantly different relative abundance among the three porous media at Day 21 and 14, respectively. No taxa were significantly associated with sampling location (i.e., top or bottom of a sampler column, an influent reservoir). These results underscore the critical influence of time on microbial taxonomic shifts and suggest that aquifer sediment is a more effective porous media material for capturing temporally dynamic taxa.

**Figure 5.**
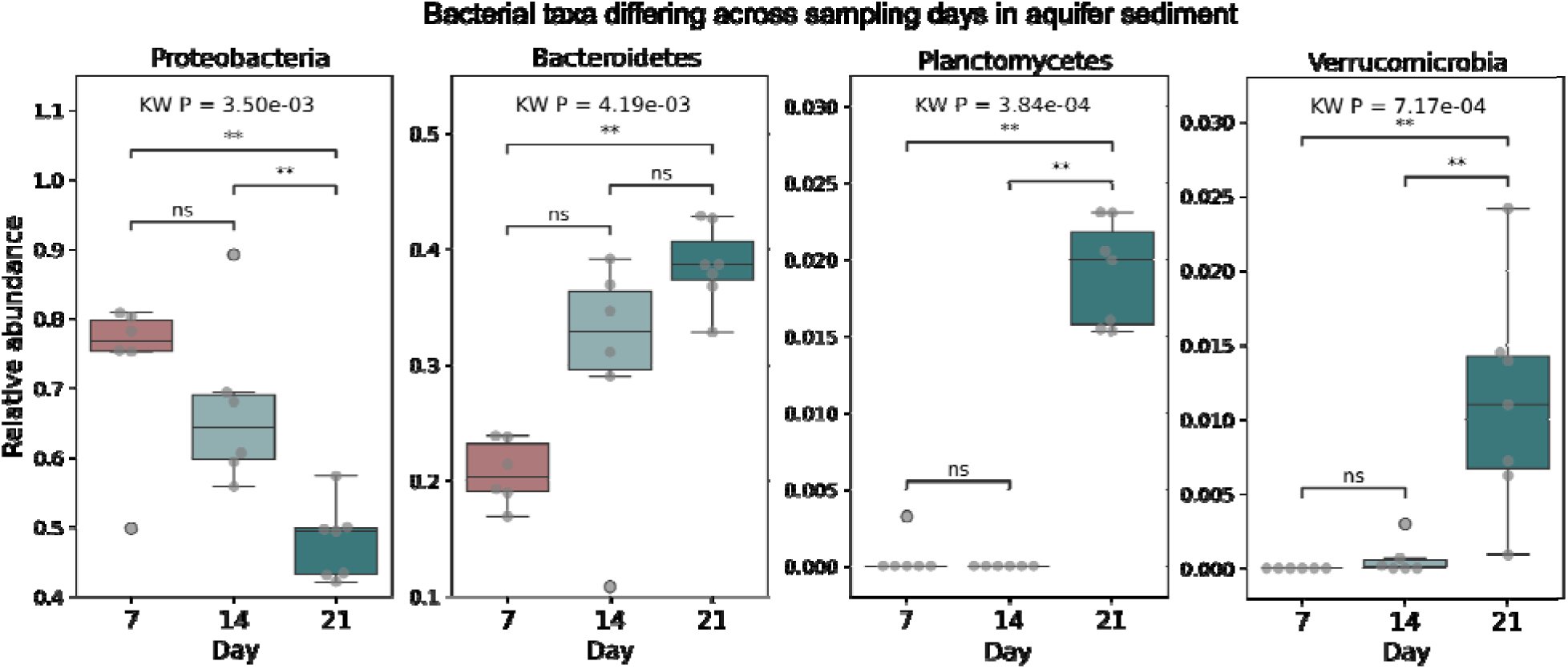
Boxplots showing four phyla with significantly different relative abundances across sampling days in aquifer sediment samples (Kruskal-Wallis [KW] P < 0.05 and adjusted P < 0.1 after Bonferroni correction). ** and ns denotes two-sided Mann-Whitney U P < 0.01 and > 0.05 (not significant), respectively. Box plots show the interquartile range (IQR), with the line representing the median and whiskers extending to 1.5 times the IQR.

## 4. Discussion

In this study, we assessed three candidate porous media - zirconia beads, native aquifer sediment and laboratory-grade silica sand - for their ability to capture temporal and spatial variation in microbial communities within a laboratory-scale passive sampler designed to mimic a sampling environment of injection of advanced treated wastewater used for managed recharge into an aquifer system. All media types reflected microbial succession over the 21-day deployment, with significantly different profiles. This result suggest that the sampling day influences the conclusions drawn, depending on the porous medium used, and underscore the importance of long-term monitoring of microbial communities in MAR and other engineered applications. It has been reported that the diversity and composition of microbial communities shift over time in aquifers and other aquatic ecosystems during recharge events based on field studies, likely attributed to prolonged exposure to nutrients like oxygen, carbon, and dissolved compounds (Yan et al., 2021 Gorski et al., 2020) and observable changes in aquifer permeability (Merino et al., 2022)(Epting et al., 2018). Accordingly, it is essential that the porous medium selected can capture the microbial community shifts resulting from these environmental changes. Among the tested porous media types, native aquifer sediment most effectively captured dynamic changes in microbial community structure of aqueous influent, in this case, advanced treated wastewater used as influent for an MAR project, suggesting its ability to resolve microbial succession and maintain community fidelity over time and its ability to study the profile of aquifer microbials communities in MAR systems.

Aquifer sediment media also exhibited greater sensitivity to environmental shifts compared to other media types, with Proteobacteria, Bacteroidetes, Planctomycetes, and Verrucomicrobia being more sensitive to temporal changes in the sampler, highlighting their potential to serve as key indicators of microbial community shifts and environmental transitions in MAR contexts. The heightened temporal sensitivity of these phyla likely stems from their functional roles in nutrient cycling, rapid adaptability, and susceptibility to redox and chemical changes that occur as water infiltrates through the aquifer system (Wang et al., 2013; Xia et al., 2022). The decline in Proteobacteria and increase in Planctomycetes, Bacteroidetes, and Verrucomicrobia over time may reflect a succession from fast-growing, metabolically flexible colonizers to slower-growing, surface-adapted specialists. Consistent with this finding, benthic bacteria such as Planctomycetes have been observed to grow more successfully on aquifer sediment, which contains different minerals with variations in size, in addition to organic carbon, than on clean silica sand (Probandt et al., 2018). Also, many of the taxa increasing over time (e.g., Planctomycetes, Verrucomicrobia) are known for biofilm formation and surface colonization, supporting the use of passive microbial samplers as mimics of sediment-associated microbial niches.

Analysis of dominant taxa at each taxonomic level across porous media types revealed the presence of microbes that are common in engineered systems, such as MAR. For example, at the class level, Gammaproteobacteria are known for their metabolic versatility and stress tolerance and dominating in nutrient-rich or disturbed environments, such as engineered aquatic systems, contaminated aquifers, and wastewater-impacted sediments (D. Li et al., 2012). Their increase in influent reservoirs of aquifer sediment over time observed in this study indicates potential adaptation to enriched organic conditions or residual contaminants in treated wastewater (J. Li et al., 2022). Betaproteobacteria are important for nitrogen cycling and include members associated with nitrification and oligotrophic growth. In this study, Betaproteobacteria were enriched at early timepoints and samples taken from the top of the sampler, consistent with higher oxygen availability. At the family level, Comamonadaceae, frequently associated with nitrate reduction and dissolved organic carbon (DOC) utilization in recharge environments (Ben Maamar et al., 2015) and presence in aquifers, were consistently abundant at early timepoints and in upper sampler regions across all porous media types. This pattern suggests their role as early colonizers in MAR systems, especially where recent recharge introduces labile organic carbon and electron acceptors such as nitrate. It may also reflect niche preference for oxic conditions typically found in samples coming from the top of the sand and zirconia media, where active nitrification-denitrification dynamics can occur. In contrast, Rhodobacteraceae exhibited greater relative abundance in influent reservoir samples, especially at later timepoints, where higher concentrations of DOC and reduced redox conditions were more likely to occur. The increase in Rhodobacteraceae over time may reflect successional shifts in microbial communities from autotrophic or oligotrophic pioneers to heterotrophic taxa. These findings align with previous MAR and groundwater studies where Rhodobacteraceae populations responded positively to high-DOC recharge, contaminant load, or geochemical stress (Chen et al., 2024; Guo et al., 2025; Yanuka-Golub et al., 2024). At the genus level, the co-occurrence of *Rhodoferax*, *Comamonas*, and *Paracoccus* across samples reflects a functionally complementary assemblage of metabolically versatile bacteria that play central roles in early colonization, nitrogen cycling, and redox transitions within recharge systems. *Comamonas*, a well-known denitrifier from the Comamonadaceae family, was consistently enriched in top sampling locations among sand and zirconia media during early timepoints, aligning with oxygenated and nitrate-rich conditions (M. Zhang et al., 2024). Similarly, *Paracoccus*, noted for its facultative aerobic denitrification and metabolic plasticity, was prominent in nitrate-influenced samples from early timepoints. In contrast, *Rhodoferax* appeared as a likely early colonizer and biofilm initiator, especially in transition zones. Its role in facultative anaerobic metabolism suggests adaptation to fluctuating redox conditions. Collectively, these taxa represent key microbial responders to hydrological and geochemical shifts in MAR environments and offer insight into niche partitioning and functional redundancy during microbial community assembly and their ability to cultivate within the bulk media of the passive sampler.

Here, we summarized the features of candidate sampling media for profiling of aquifer microbial communities, including microbial characteristics and temporal dynamics in addition to aquifer structural complexity, porosity, and permeability (**Table 1**). Overall, native aquifer sediment provides the most realistic and informative substrate outperforming artificial porous media (i.e., zirconia beads and laboratory-grade silica sand) in several ways. First, native sediments inherently reflect indigenous microbial composition in the aquifer, supporting high diversity across bacteria, archaea, and other microorganisms due to their natural heterogeneity (Castelle et al., 2013; Flynn et al., 2013; Hug et al., 2015). The complex physical structure of aquifer sediments, including micropores, varied mineral surfaces, and differing grain sizes, facilitates microbial attachment, niche differentiation, and the development of intricate community structures (Barba et al., 2019; Li et al., 2013). These structural features also allow aquifer sediments to capture natural temporal dynamics, reflecting seasonal or environmental shifts in microbial populations (Abiriga et al., 2021; Tian et al., 2022), and likely make these sediments less sensitive to sampling locations when used in a passive sampler. Additionally, the porosity and permeability of native sediments influence nutrient and oxygen fluxes, further supporting diverse and functionally relevant microbial assemblages (Abiriga et al., 2021; Chang et al., 2023). In contrast, zirconia beads and silica sand lack the environmental complexity to accurately mimic natural microbial habitats, offering simplified communities and limited temporal variation (Miao et al., 2021). Furthermore, aquifer sediment and silica sand were easy to use during the bench-scale experiment, whereas zirconia beads proved problematic due to clogging, likely caused by the decreased void space and biofilm growth within the cycling reservoir, which led to the collapse of media columns during operation and reduced the feasibility of their use. From a practical standpoint, native aquifer sediment offers representative microbial and geochemical conditions that closely mimic field infiltration and recharge behavior, whereas lab-grade silica sand provides uniform grain size and chemistry for experimental consistency, and zirconia beads, though commercially available, were found to be prone to swelling and bursting under operational conditions by us and others (Fakhreddine et al., 2021; Kim et al., 2010). Collectively, these characteristics indicate that native aquifer sediments are superior for representing microbial community diversity, structure, temporal shifts, and ecological interactions *in situ*, making them the optimal sampler medium for monitoring microbial communities in MAR and other groundwater applications.

**Table 1.** Summary of comparative characteristics of porous media used in this study.

| Feature | Native aquifer sediment | Zirconia beads | Lab-grade silica sand |
| --- | --- | --- | --- |
| **Microbial Characteristics** | Reflects indigenous microbial communities; supports high diversity and complex community structures due to natural heterogeneity | Limited support for natural communities; simplified structures | Supports microbial growth but simplified community structures; diversity influenced by surface properties |
| **Temporal Dynamics** | Captures natural seasonal and environmental shifts in microbial populations | Limited; artificial conditions restrict temporal variation | Moderate shifts; may not fully reflect natural dynamics |
| **Structural Complexity** | Micropores, varied mineral surfaces, heterogeneous grain sizes; facilitates attachment and niche differentiation | Uniform surface; lacks micropores and mineral variety | Limited surface heterogeneity; smoother grain surfaces |
| **Porosity & Permeability** | Natural porosity supports nutrient and oxygen fluxes, sustaining diverse and functionally relevant microbial assemblages | Minimal; does not mimic natural fluxes | Moderate; influences nutrient diffusion but simplified |
| **Practical considerations** | Representative microbial and geochemical conditions, better mimic field infiltration and recharge behavior. | Commercially available, prone to swelling and bursting | Uniform grain size and chemistry minimize variability |

## 5. Conclusions

This study shows that an aquifer sediment-based passive sampler is an effective tool for tracking changes in aquifer microbial communities over time. Due to the significant influence of time on microbial community structure, it is critical to longitudinally monitor microbial communities in engineered systems such as managed aquifer recharge. To account for temporal shifts in microbial succession, including the attenuation of notable biofilm forming taxa such as Planctomycetes and Verrucomicrobia, it is recommended that samples be collected in parallel after the same acclimation period to enable valid comparisons across conditions within a study. Also, the selection of porous media for samplers is critical, as they differ in their capacities in capturing microbial communities. Native aquifer sediment outperformed laboratory-grade silica sand and zirconia beads for better capturing microbial community dynamics reflective of the influent water, as indicated by its microbial community structure uniformity across sampling locations and its rapid response to temporal changes. These advantages potentially enable the capture of microbial hazards (e.g., pathogens and antimicrobial resistance genes) and those involved in contaminant degradation over time. Silica sand performed moderately well, but lacked the microbial retention and representative community structure observed in sediment. Zirconia beads were prone to clogging due to low void space and film buildup, detracting from the feasibility of their use. Given the short time frame of this experiment (21 days) and the absence of a plateau in taxonomic abundances or stable community resemblance over the extended sampling window, it is difficult to definitively recommend definitively recommend an optimal sampling medium for long-term MAR applications. Nevertheless, for short-term sampling, our results provide practical guidance on porous-media selection for passive sampling of solid-phase–associated microbial communities, contributing to the development of a customizable sampler for *in situ* microbial monitoring in subsurface environments during managed aquifer recharge and other groundwater applications.

## Acknowledgements

We are grateful to the members of LEAPH (the Liao laboratory) and the Pruden Lab for their enriching discussions. We also thank Arba Williams for finalizing the 3D print designs for printing efficiency. This research was funded in part by the U.S. Environmental Protection Agency (No. RD-84061901), the Hampton Roads Sanitation District, the National Science Foundation NRT Award 2125758, the Virginia Tech College of Engineering, and the Virginia Tech College of Engineering New Horizons Program.

## Data availability

16S rRNA gene amplicon sequencing reads have been deposited at the NCBI Sequence Read Archive (SRA) under the BioProject number PRJNA1353100.

## Author contributions

JL and AP designed the study. MR and KT conducted the experiments. MR, HB, and JL analyzed the data. MR and HB wrote the paper with input from JL, AP, and MS.

