## Supplemental figures and tables for "Assessment of a Passive Sampler for Aquifer Microbial Community Profiling and Comparison of Porous Media Applied in a Managed Recharge Scenario"

**Supplementary figures**

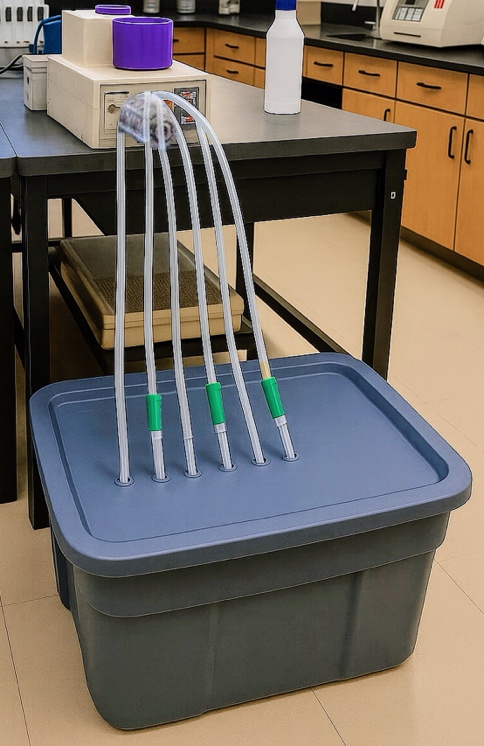

**Figure S1**. Setup of bench-scale experiment for investigating passive microbial samplers. A 20-L reservoir of SWIFT effluent is pumped continuously through samplers containing the candidate media. Three media were tested in triplicate: aquifer sediment, sand, and zirconia beads. The media were packed in green 3D printed columns to stabilize the suspended media and fed from independent reservoirs in a continuous loop, using the same pump head to achieve relatively consistent flow rates through the columns.

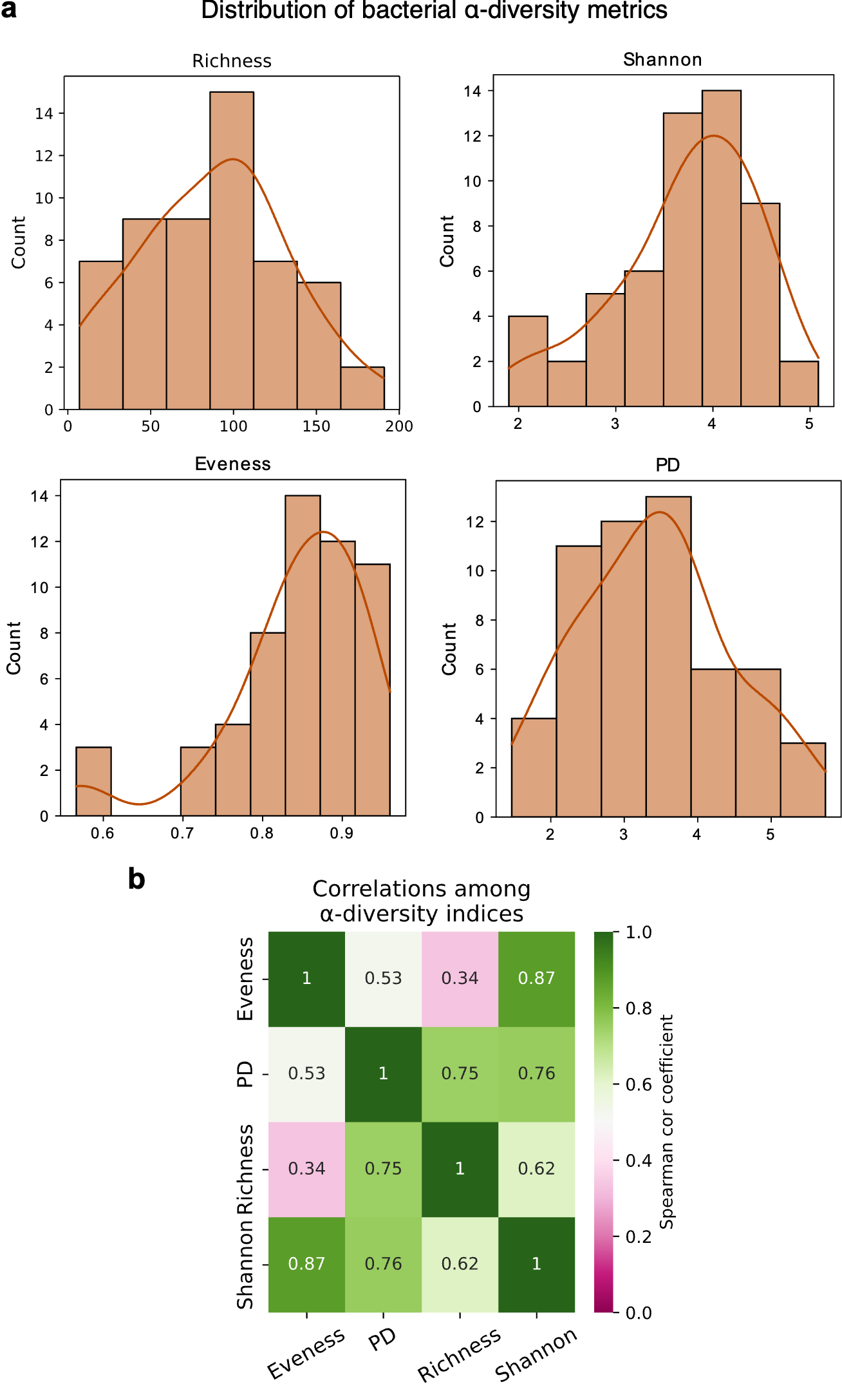

**Figure S2**. Distribution and correlations of α-diversity metrics. (a) Histograms showing the distribution of α-diversity metrics, including Richness, Shannon-Wiener diversity index, Simpson’s evenness, and Faith’s phylogenetic diversity (PD). (b) Heatmap illustrating the pairwise Spearman's rank correlation between α-diversity metrics.

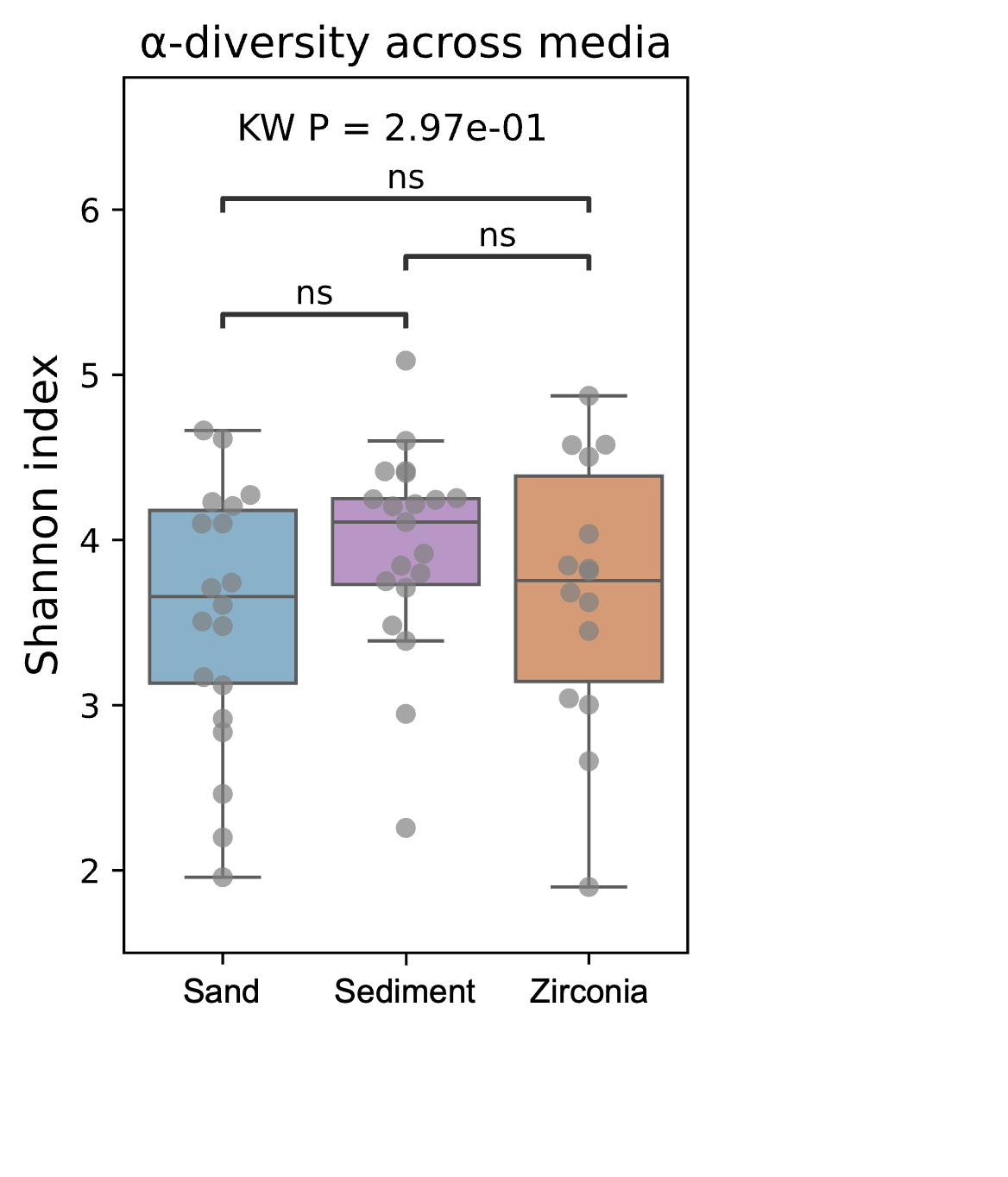

**Figure S3**. α-diversity of microbial communities compared across three different media (sand, aquifer sediment, and zirconia beads). α-diversity is represented by the Shannon-Wiener index. KW: Kruskal–Wallis test. "ns" denotes “not significant” in the pairwise comparisons.

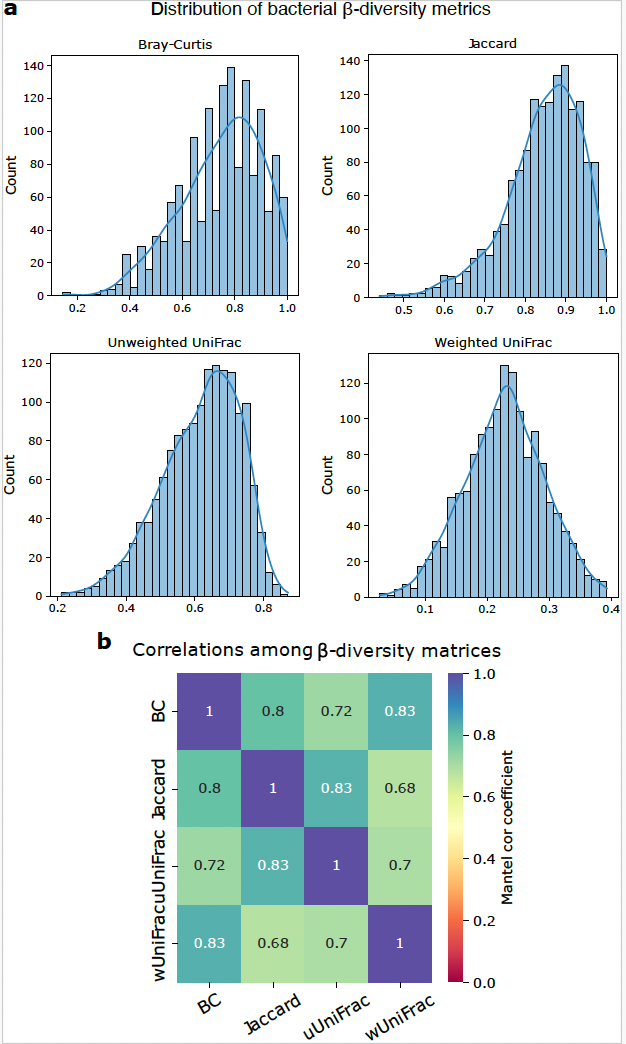

**Figure S4**. Distribution and correlations of β-diversity metrics. (a) Histograms showing the distribution of β-diversity metrics, including Bray-Curtis (BC), Jaccard, unweighted UniFrac (uUniFrac), and weighted UniFrac (wUniFrac) distances. (b) Heatmap illustrating the pairwise Mantel correlation between β-diversity metrics.

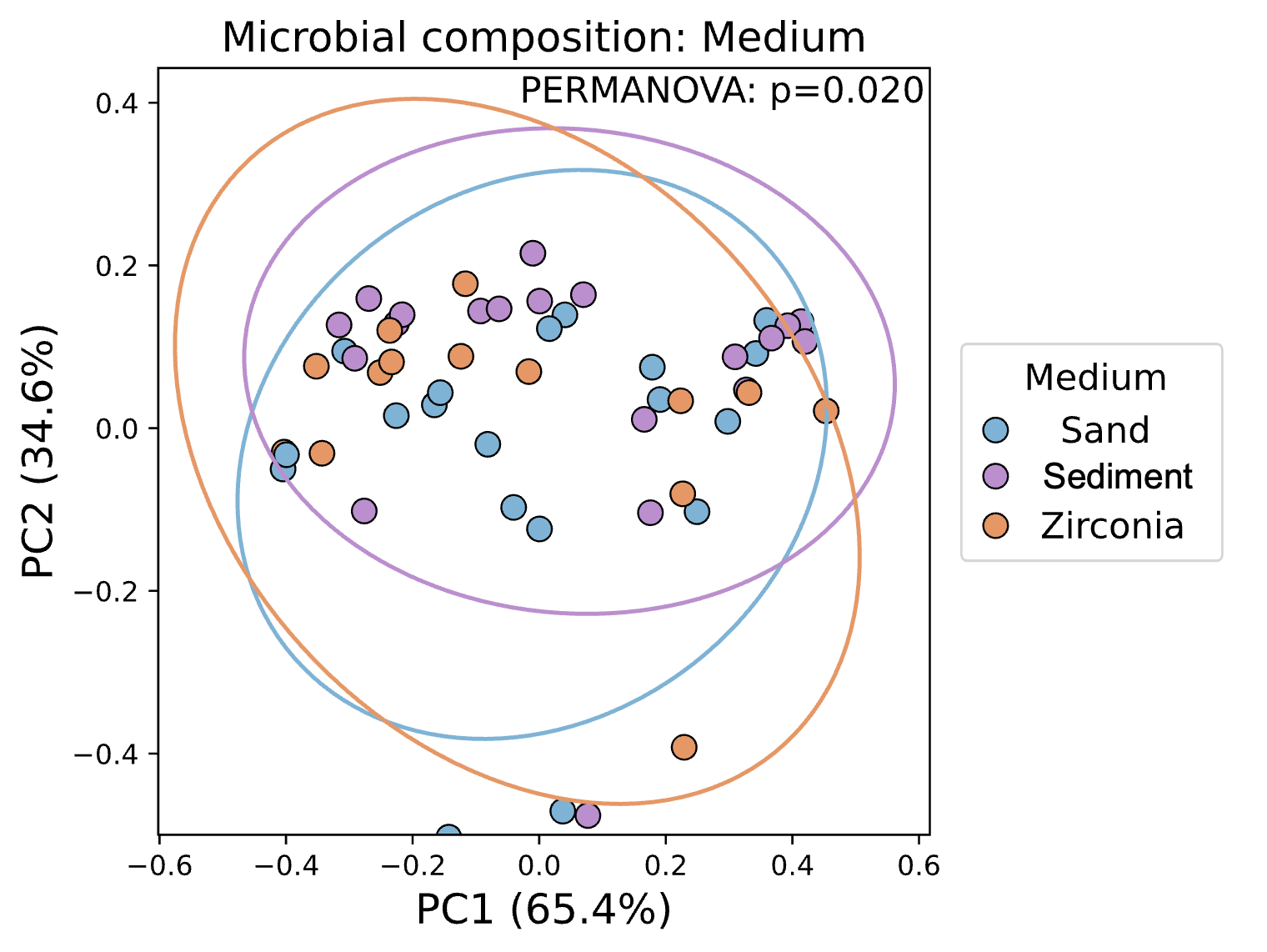

**Figure S5**. Principal coordinate analysis (PCoA) of microbial community composition based on Bray–Curtis dissimilarity. Dots are colored by medium type (sand, aquifer sediment, zirconia). PC1 (Principal Component 1) and PC2 (Principal Component 2) is the first and second principal coordinate, respectively. The percentage indicates the proportion of the total variance in the data that is captured by that specific axis. PERMANOVA: Permutational Multivariate Analysis of Variance.

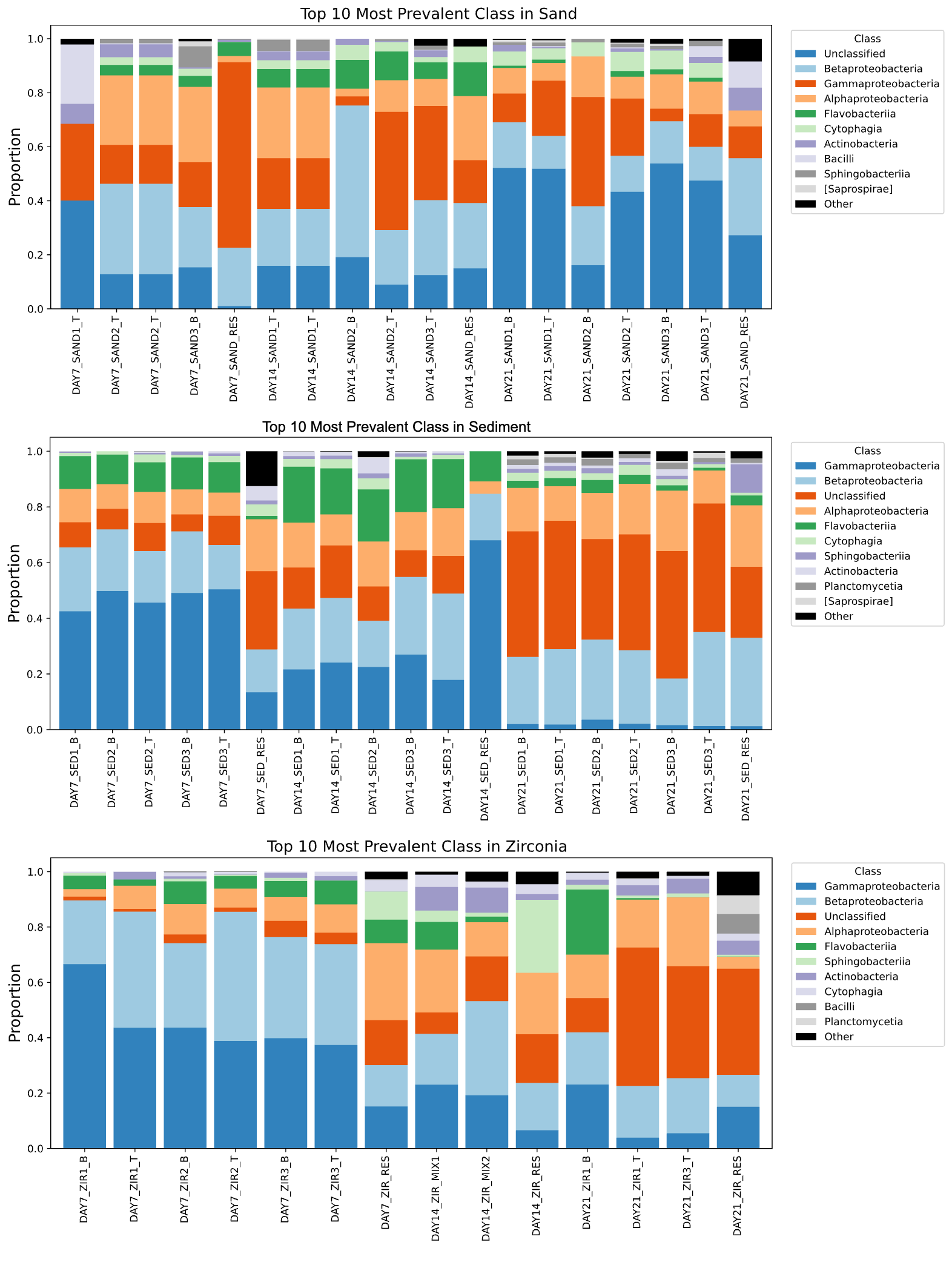

**Figure S6**. Stacked bar plots showing the relative abundance of the top 10 most prevalent classes across samples of each porous medium (sand, aquifer sediment, and zirconia beads)

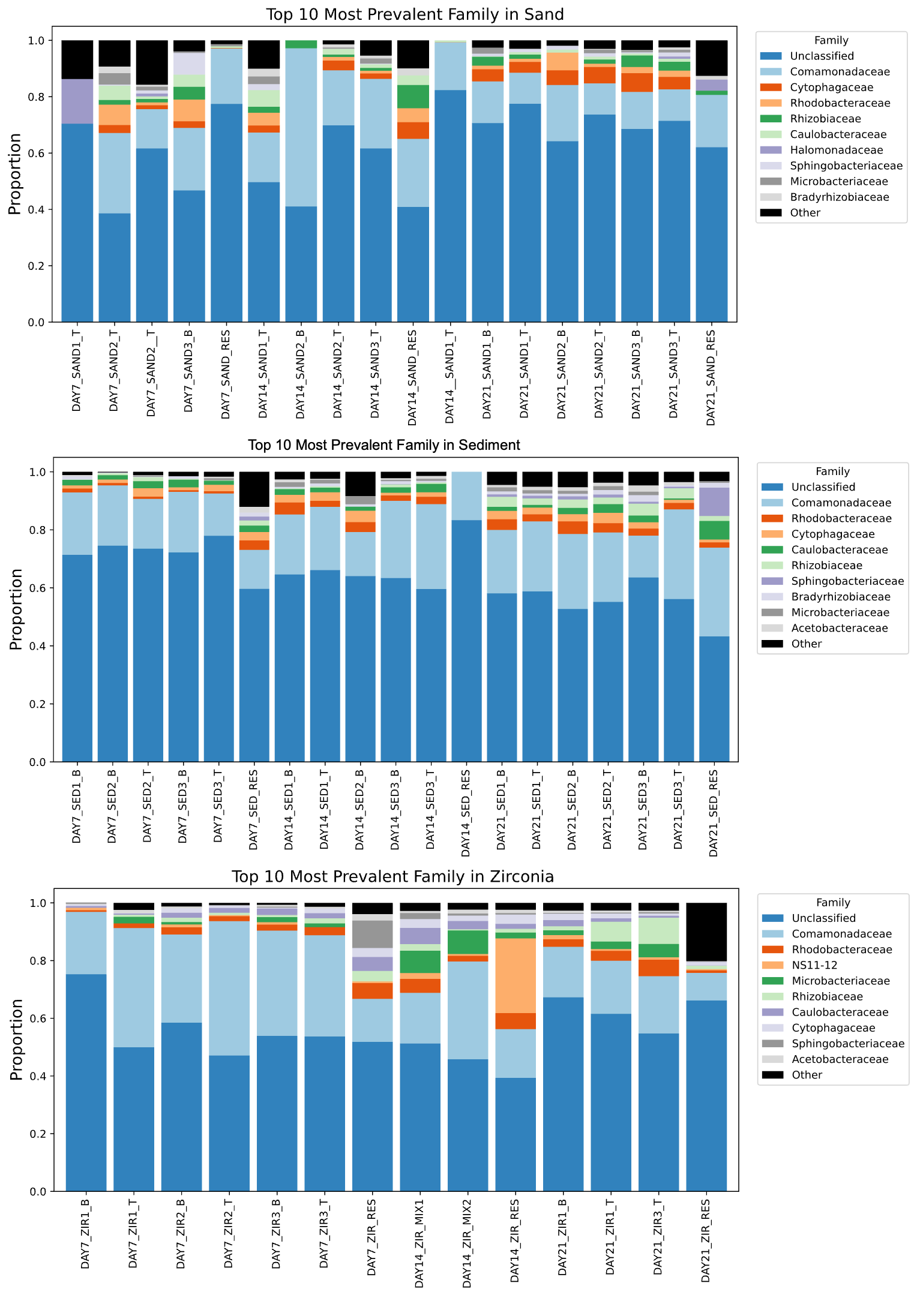

**Figure S7**. Stacked bar plots showing the relative abundance of the top 10 most prevalent families across samples of each porous medium (sand, aquifer sediment, and zirconia beads).

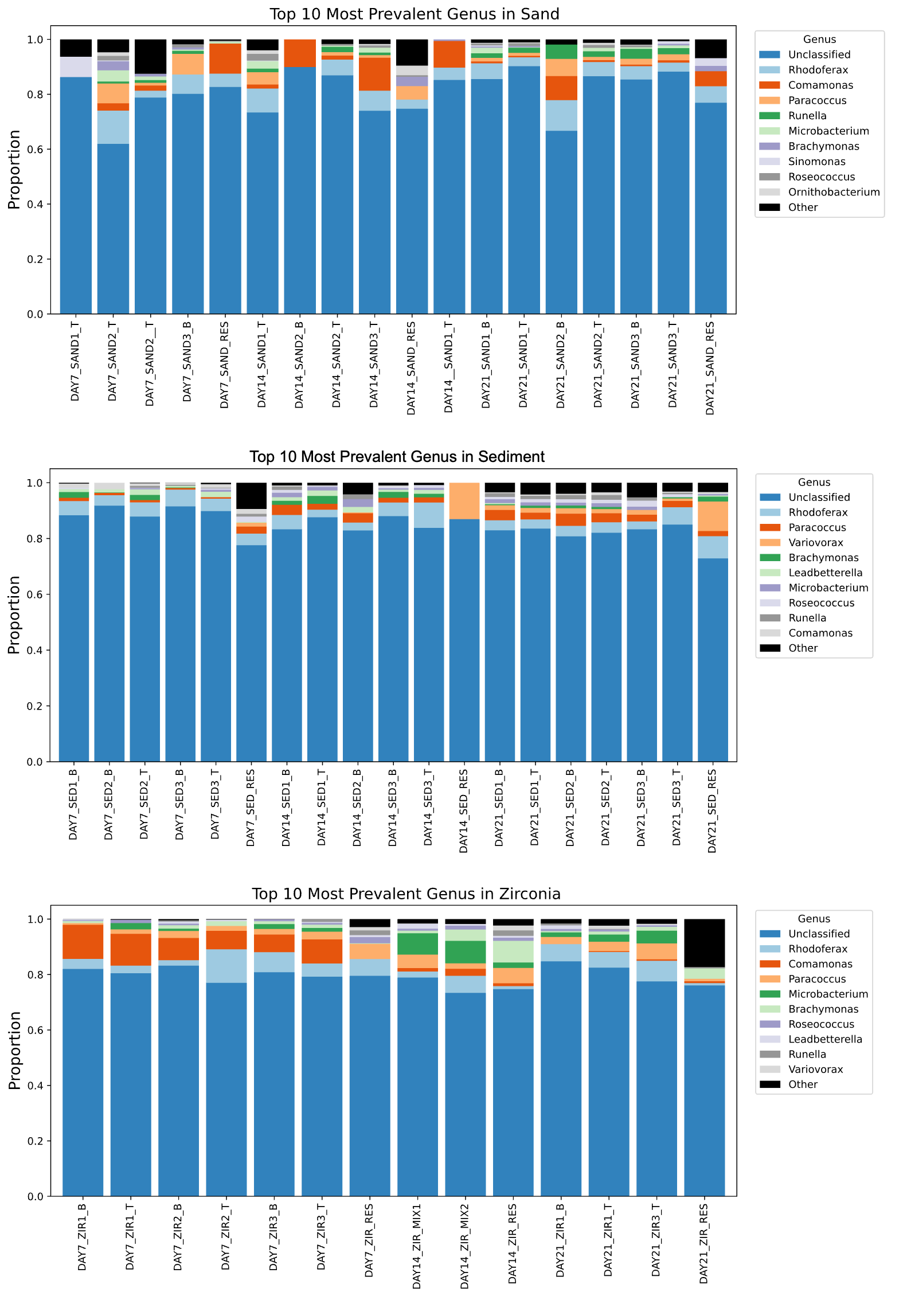

**Figure S8**. Stacked bar plots showing the relative abundance of the top 10 most prevalent genus across samples of each porous medium (sand, aquifer sediment, and zirconia beads).

**Supplementary tables**

**Table** **S1**. Bacterial taxa with significantly different relative abundances across comparison conditions.

| **Taxa Name** | **Comparison Condition** | **H Statistic** | ***P*-value** | **Adjusted *P*-value** |
| --- | --- | --- | --- | --- |
| Class | | | | |
| gammaproteobacteria | Day across Medium = Aquifer sediment | 13.12 | 0.00142 | 0.0667 |
| flavobacteriia | Day across Medium = Aquifer sediment | 12.67 | 0.00178 | 0.0834 |
| bacteroidetes unknown | Day across Medium = Aquifer sediment | 13.16 | 0.00142 | 0.0667 |
| planctomycetia | Day across Medium = Aquifer sediment | *16.82* | 0.000223 | 0.0105 |
| [pedosphaerae] | Medium across Day = 21 | 12.67 | 0.00177 | 0.0833 |
| [pedosphaerae] | Day across Medium = Aquifer sediment | 13.60 | 0.00111 | 0.0524 |
| Order | | | | |
| ellin329 | Day across Medium = Aquifer sediment | 14.08 | 0.000875 | 0.0788 |
| pirellulales | Day across Medium = Aquifer sediment | 16.82 | 0.000223 | 0.0204 |
| opitutales | Medium across Day = 14 | 13.80 | 0.00101 | 0.0909 |
| Family | | | | |
| pirellulaceae | Day across Medium = Aquifer sediment | 16.82 | 0.000223 | 0.0296 |
| Genus | | | | |
| *A17* | Day across Medium = Aquifer sediment | 16.82 | 0.000223 | 0.0363 |
| Species | | | | |
| *planctomycetes species unknown* | Day across Medium = Aquifer sediment | 16.82 | 0.000223 | 0.0372 |
